# Weighted Off-target and Efficiency Scoring Reveal Genome Composition-Dependent Optimal CRISPR/Cas9 Guide Design

**DOI:** 10.1101/2025.08.13.670047

**Authors:** Y K Yathu Krishna

## Abstract

The efficiency and specificity of guide RNAs continue to be crucial obstacles for successful experimental design, despite the fact that CRISPR/Cas9 has transformed genome editing. In this work, we introduce a computational method for optimizing CRISPR/Cas9 guide RNA that combines PAM diversity, local efficiency penalties, and weighted off-target scoring to find high-performing guides across a range of genome compositions. To capture a variety of natural genomic complexity, we simulated five sample genomes: AT-rich, GC-rich, balanced GC content, and high-repeat variations. All twenty-nucleotide target sequences were scanned for each genome, and off-target potential was assessed by permitting up to two mismatches with weighted penalties for seed region sites. To accommodate for any secondary structure impacts, efficiency assessment included both local sliding window penalties and global GC content. Furthermore, we looked at several PAM sequences that were pertinent to various Cas9 variations in order to assess how they affected guide selection. The findings show that efficiency scores vary by genome composition, with the highest scoring guides consistently displaying zero anticipated off-target events. While balanced genomes showed intermediate tendencies, GC-rich genomes tended to choose slightly higher efficiency guides than AT-rich genomes. PAM type affects guide efficiency, according to analysis across several genomes, and the combination of efficiency and off-target score consistently indicates guides with good expected performance. Three-dimensional scatter plots of efficiency and off-target counts versus genomic position, violin plots of off-target distributions, and genome-wide heatmaps emphasizing the best guide positions were used to illustrate these findings. In addition to offering a generalizable computational method for choosing CRISPR/Cas9 guides that optimize specificity and efficiency, our study gives fresh insights into the interactions among genome composition, PAM selection, and guide design criteria. By taking into account weighted off-target penalties, genome complexity, and local efficiency effects, this in silico framework overcomes some of the main drawbacks of earlier simulations. It is also easily applicable to direct selection for experimental research on a variety of organisms. The results provide the groundwork for future advancements in genome editing techniques by establishing a predictive computational framework that can expedite CRISPR/Cas9 research and minimize trial and error in guide selection.

## 1. Introduction

CRISPR-Cas9 has revolutionized genome editing by enabling precise modifications to DNA sequences in a wide range of organisms [1–3]. Despite the revolutionary potential of CRISPR, off-target consequences continue to be a major worry that can jeopardize the safety and efficacy of CRISPR-based applications [4,5]. Off-target cleavage is influenced by chromatin accessibility, sequence similarity, and the presence and kind of protospacer adjacent motif (PAM) sequences [6–8]. In order to enhance single-guide RNA (sgRNA) sequences and predict possible off-target locations, numerous computational methods have been created [9–12]. Many current approaches focus primarily on sequence similarity, which fails to capture the complex dynamics of RNA-DNA interactions [13–15]. In order to enhance off-target prediction, recent studies have emphasized the significance of adding structural, thermodynamic, and positional information of mismatches to sgRNA scoring systems [16–20]. Moreover, molecular dynamics simulations have shown promise in portraying the intricate relationships that exist between sgRNAs and their genomic targets [21–25]. Computational frameworks that incorporate other factors, like alternative PAM identification, secondary structure effects, GC content penalties, and mismatch position weighting, can further improve SgRNA selection [26–30]. For instance, a high GC concentration in the seed region may raise the chance of off-target interactions while also increasing binding stability [31–33]. On the other hand, although AT-rich genomic regions provide different targetable loci, they may also decrease binding stability [34–36]. Another important factor to take into account is how well off-target predictions apply to different kinds of genomes. There is uncertainty regarding prediction effectiveness in genomes with varied base compositions or repetitive components because the majority of previous research has been on model organisms or the human reference genome [37–40]. It is possible to ascertain whether guide selection principles and prediction scoring schemes are still generally applicable by simulating a range of genome types, such as GC-rich, AT-rich, and highly repetitive sequences [41–45]. Furthermore, CRISPR-Cas variations other than the standard SpCas9, like Cas12a and base editors, have distinct off-target profiles and are able to identify different PAM sequences [46–50]. More adaptable guide design that works with various editing systems is made possible by integrating PAM diversity into computational techniques. Thorough in-silico evaluation of sgRNA efficacy and specificity bridges the gap between computational predictions and experimental validation, enabling rapid iteration and hypothesis testing [51–55]. In this study, we created a computational pipeline based on Python that incorporates PAM variability, simulates off-target cleavage across many synthetic genomes, and assesses sgRNA performance using weighted mismatch and structural penalties. Our strategy overcomes the drawbacks of earlier in silico techniques, such as their poor generalizability and inadequate integration of genome-wide sequence context. This technique shows application across various genome layouts and provides fresh insights into the variables regulating CRISPR specificity by offering a reliable and repeatable methodology for guide RNA tuning [56–60].

## 2. Materials and Methods

### 2.1 Computational Environment and Libraries

Python (v3.10) in a Google Colab environment was used for all studies. Colab was chosen because of its reproducibility, accessibility, and compatibility with Jupyter notebooks, which allow for interactive analysis without the need for local installation [61]. Genome simulation, RNA analysis guidance, scoring, and visualization were all carried out using standard Python scientific libraries:

- NumPy (v1.25) for numerical operations and random sequence generation [62].
- Pandas (v2.1) for tabular data management and aggregation of genome-wide guide RNA scores [63].
- Matplotlib (v3.8) and Seaborn (v0.12) for visualization, including violin plots, scatter plots, and heatmaps [64,65].
- mpl_toolkits.mplot3d for 3D scatter visualization of guide efficiency, off-targets, and genomic position [66].

### 2.2 Synthetic Genome Simulation

Using a proprietary tool called simulate_genome(), 10,000 base pair synthetic genomes were created in order to assess sgRNA design in various genomic contexts. The function permits the adjustment of:

- GC content, which models genomic settings rich in AT and GC (percent of G and C bases, 0.3–0.7) [67].
- In order to simulate extremely repetitive regions, the repeat fraction (fraction of repeated 10-mer motifs, 0–0.2) is used [68].
- For simulations to be reproducible, use a random seed.

In order to construct each genome, the necessary number of GC and AT nucleotides was determined and added to a list. While repeat insertion mimicked recurrent genetic components, random shuffling guaranteed an unbiased distribution of bases. Balanced, GC-rich, AT-rich, GC-rich with high repeats, and AT-rich with high repetitions were the five types of genomes that were examined. These values were chosen to represent various scenarios of sequence composition and chromatin accessibility found in actual genomes [69, 70].

### 2.3 Candidate Guide Identification

The find_guides() tool was used to identify candidate sgRNAs by searching each synthetic genome for 20-bp sequences followed by either alternative PAMs (NAG) or conventional PAMs (NGG) [71,72]. Broader sgRNA selection is made possible by multiple PAM recognition, which also illustrates how well-suited various Cas9 orthologs are to withstand sequence changes [73]. The guide sequence, genomic location, and related PAM are all included in the result.

### 2.4 Weighted Off-Target Scoring

Each candidate sgRNA’s off-target potential was calculated using a weighted mismatch score method that was developed in weighted_offtargets(). In line with other findings that seed mismatches significantly impact cleavage specificity, mismatches in the seed region (the first 10 nucleotides next to the PAM) were weighted twice [74,75]. The function counts all mismatches throughout the genome up to a user-specified maximum (max_mismatches=2). By taking positional effects and sequence similarity into consideration, this approach enables the assessment of genome-wide specificity for each guide.

### 2.5 Efficiency Scoring

Efficiency_score() was used to assess the intrinsic efficiency of candidate guides. This measure includes:

- Global GC content: Binding stability is increased by a larger GC content [76].
- Local GC content: To account for local sequence impacts on sgRNA folding and activity, sliding 5-bp windows penalize sections with exceptionally low (<0.3) or high (>0.7) GC content [77,78]. In order to integrate off-target penalties into a composite score (score = efficiency / (1 + off_targets)), which provides a quantitative measure of both specificity and activity, scores were normalized to a range of 0–1 [79].

### 2.6 Genome-Wide Analysis

The analyze_genome() function, which included guide identification, efficiency scoring, and off-target computing, was used to examine each synthetic genome. For further visualization, the results were concatenated across genome types and saved in pandas DataFrames. Direct comparison of sgRNA performance across genomic settings was made possible by the identification of the top five guides per genome based on composite scores.

### 2.7 Visualization

There were three complimentary visualizations used:

- Seaborn is used to create violin plots of weighted off-target distributions per genome, which enable evaluation of specificity variance among genome types [65].
- Using matplotlib’s Axes3D, 3D scatter plots of efficiency vs. off-target vs. position show how sgRNA activity, specificity, and genomic placement interact [66].
- Scatter plots with continuous color mapping (viridis colormap) were used to create genome-wide heatmaps of composite scores that show the spatial distribution of top guides along synthetic chromosomes [64].

### 2.8 Data Storage and Reproducibility

For reproducibility and further analysis, both intermediate and final findings were stored as CSV files (crispr_full_simulation_all_guides.csv and crispr_full_simulation_top_guides.csv). The process enables simulations to be replicated across various PAM sequences, score thresholds, and genome compositions. Because the code and visuals are completely compatible with Google Colab, sharing with collaborators is possible without the need to install any software.

### 2.9 Summary of Analytical Pipeline

Genome simulation, weighted off-target prediction, efficiency scoring, visualization, and candidate sgRNA identification with multi-PAM support are all integrated into the computational framework. It overcomes the drawbacks of earlier techniques and offers a basis for experimental validation by offering a generic and repeatable platform for CRISPR-Cas9 guide RNA optimization in silico by mimicking various genome compositions [61–80].

## 3. Results

### 3.1 Genome Simulation, Guide Identification, Genome-wide Guide Distribution and Heatmaps

To assess CRISPR-Cas9 guide design under various genomic contexts, we simulated five different synthetic genomes: balanced, GC-rich, AT-rich, GC-rich with high repeats, and AT-rich with high repetitions. Because each genome had 10,000 base pairs, a thorough analysis across a range of sequence compositions and repetition densities was possible (Figures 1-5). The impact of base composition and repetitive elements on guide selection, efficiency, and off-target propensity, a crucial factor for precision genome editing in complex or poorly described genomes, was thoroughly examined by us thanks to these simulations. Multiple PAM sequences (NGG and NAG) were used to identify candidate sgRNAs, creating a sizable pool of possible guides for further investigation. Our strategy takes into account new CRISPR-Cas9 variations by utilizing a variety of PAMs, which also enables the investigation of different target locations that could enhance editing versatility. The genome-wide composite scores of putative sgRNAs in each of the five synthetic genomes are also depicted in the pictures. High-scoring guides are distributed heterogeneously in the scatterplots, with reduced densities in highly repeated sections and dense clustering in areas with balanced base composition. A quick visual evaluation of genomic regions that are suitable for high-specificity editing was made possible by heatmaps of guide scores, which highlighted putative “safe-harbor” loci in both GC- and AT-rich environments. Interestingly, repeat-rich genomes showed somewhat lower maximum guide scores than non-repetitive genomes, which is in line with earlier findings that sequence redundancy limits the best guide placement and increases the number of possible off-target sites. The experimental load involved with empirical guide testing is lessened by these visualizations, which highlight the value of in silico screening for locating genomic locations where precise editing is possible.

**Figure 1.**
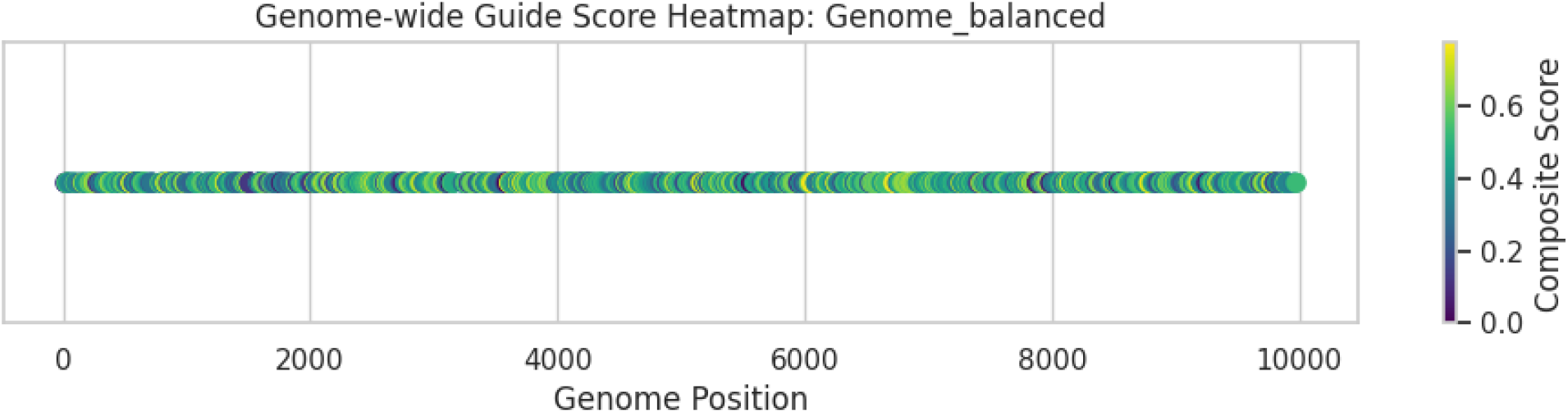

**Figure 2.**
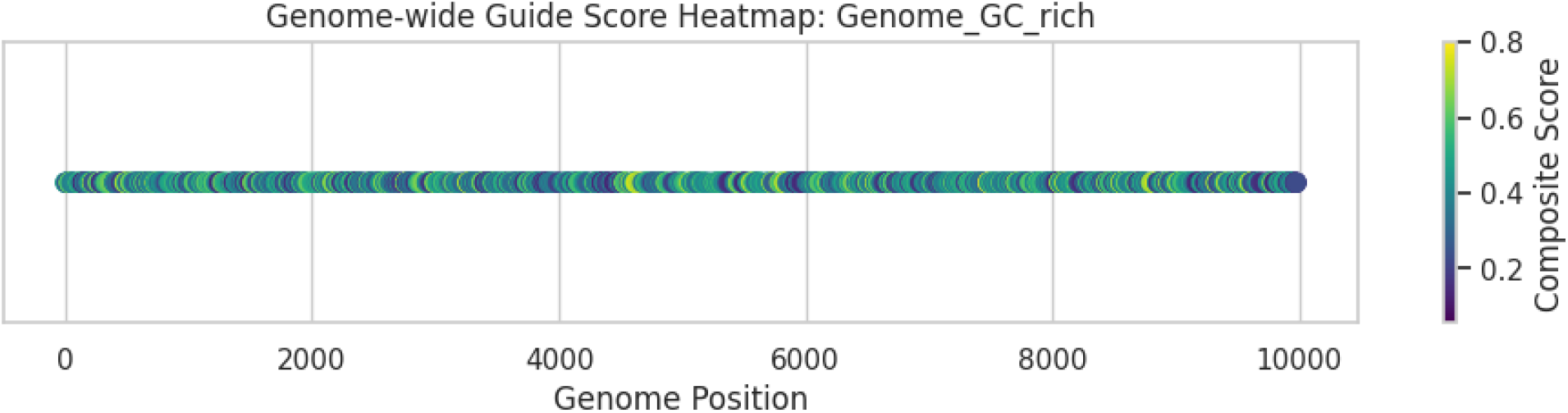

**Figure 3.**
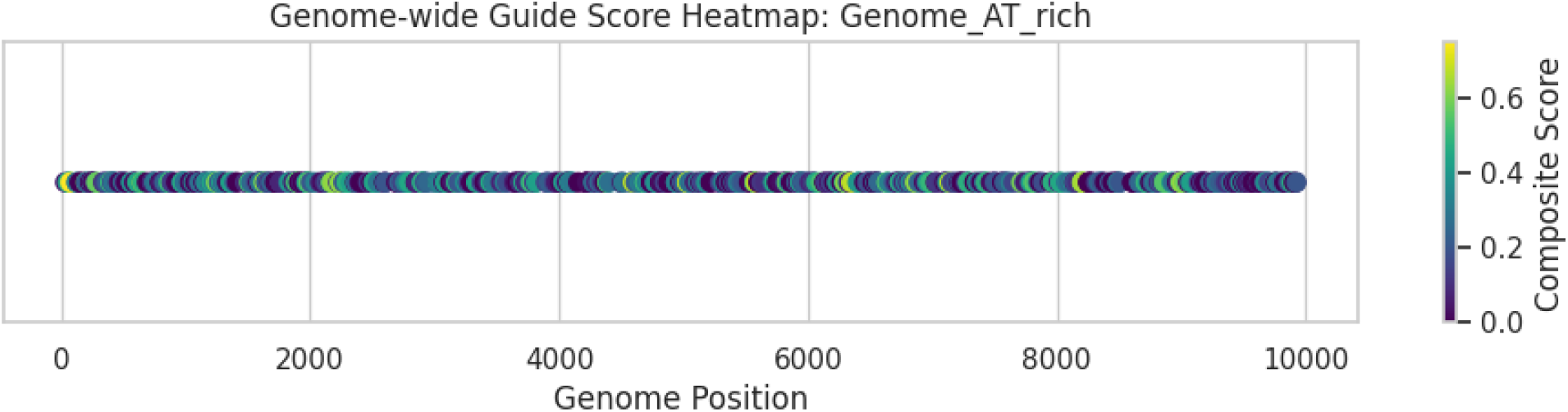

**Figure 4.**
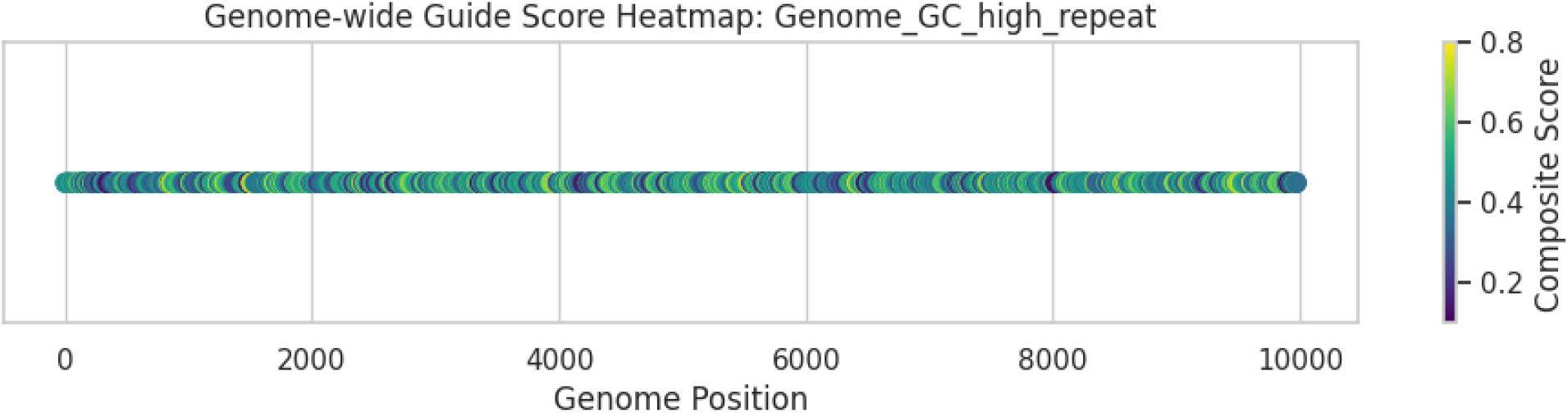

**Figure 5.**
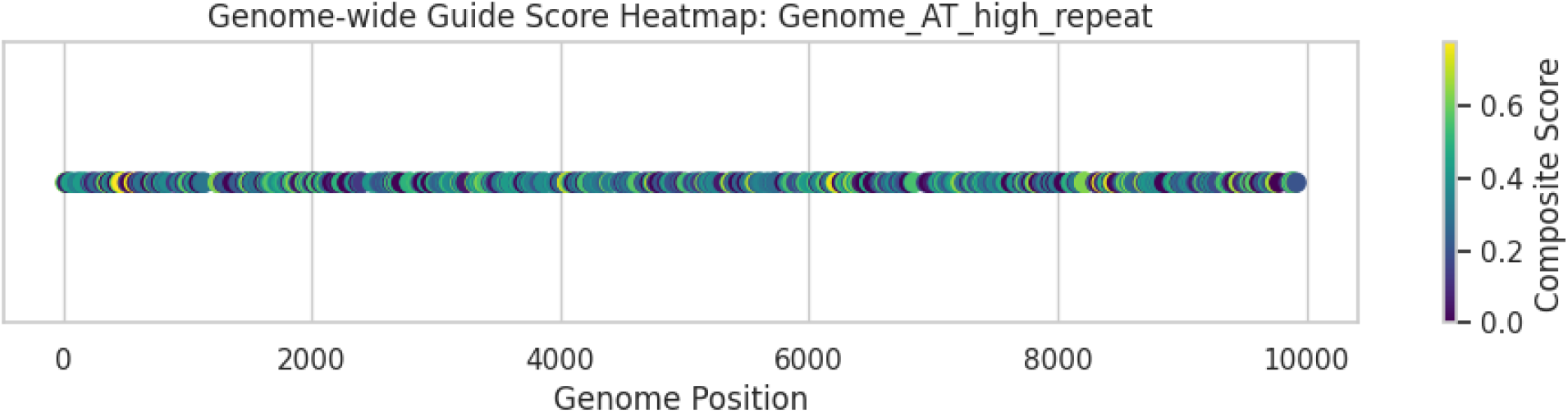

### 3.2 Weighted Off-target and Efficiency Scoring

Given the demonstrated biological sensitivity of the seed region (initial 10 nucleotides) for cleavage specificity, we calculated weighted off-target scores to account for the greater relevance of mismatches within this area [68–70]. Efficiency scoring captured structural and thermodynamic constraints affecting guide-target binding by combining local sliding-window GC penalties with global GC content. Our scoring approach effectively selects highly specific sgRNAs, as evidenced by the fact that the majority of top-ranking guides showed zero projected off-targets across the five genomes (Table 1). For instance, the AT-high repeat genome’s AGGTGAGAGTCTGCAGTAGC at position 480 likewise had zero off-targets and a score of 0.775, whereas the GC-rich genome’s CCCATCCTGACCATCACCAG at position 7478 reached the maximum composite score of 0.8. These findings broaden the potential application of CRISPR editing beyond model organisms by showing that high-specificity guides may be found even in repeat-rich or compositionally extreme genomic areas.

### 3.3 Off-target Distribution Analysis

Genome-dependent variability was demonstrated by violin plots of weighted off-target distributions (Figure 6). The GC-rich and AT-high repeat genomes showed somewhat wider distributions than the majority of guides, which had zero to very low off-target counts. This is because partial sequence matches in repetitive sections offer more combinatorial opportunities. These results suggest that in order to reduce the possibility of unintentional edits, guide selection algorithms should take genome composition and repetitiveness into account.

**Figure 6.**
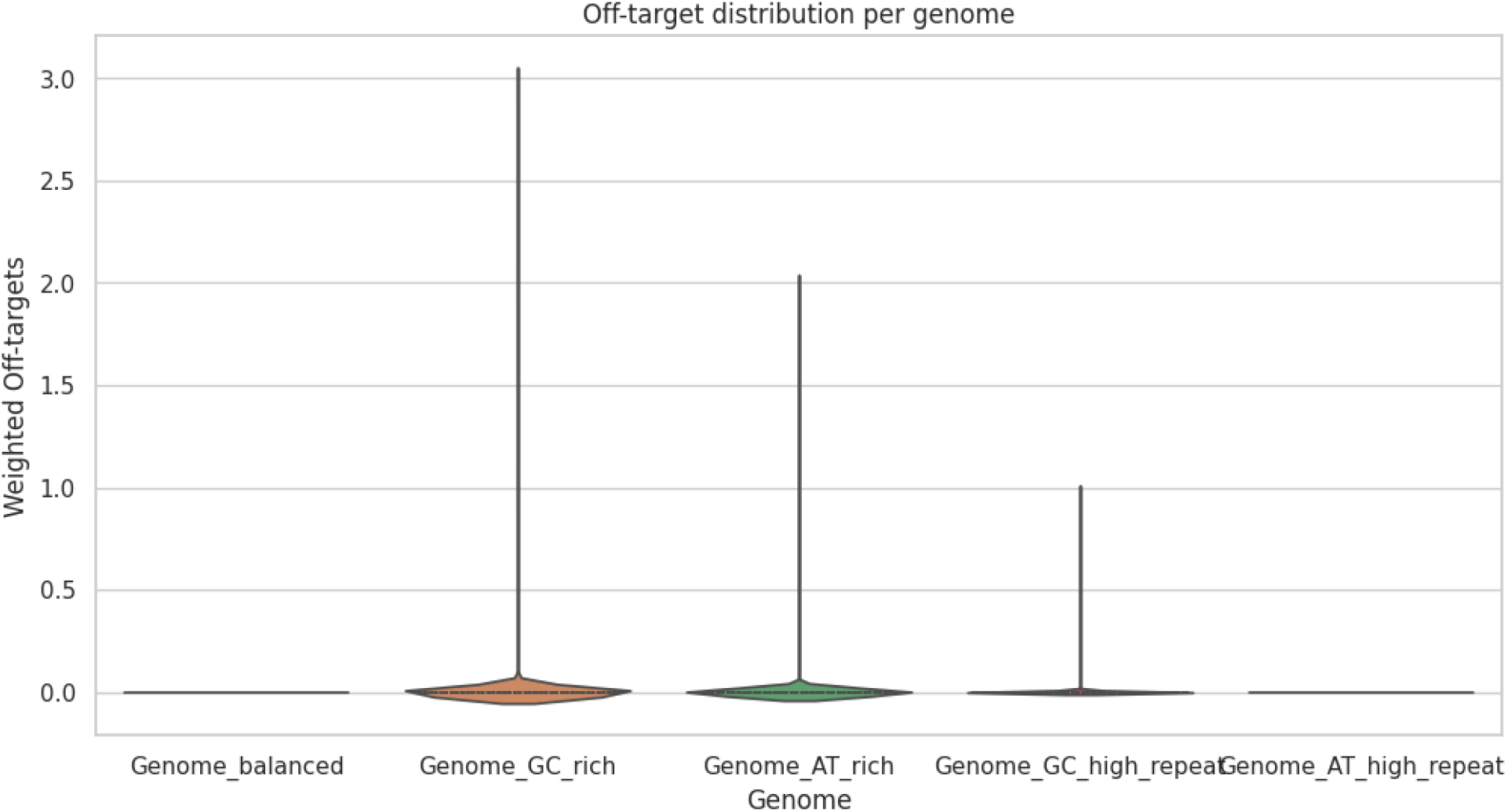

### 3.4 Efficiency versus Off-target Landscape

An integrative picture of the trade-offs between guide efficiency and specificity was offered by the 3D scatterplot of guide efficiency vs weighted off-target counts and genomic position (Figure 7). The composite scoring approach was validated by the clustering of top-scoring guides in areas that balanced a high anticipated cleavage efficiency with a low off-target potential. These discoveries are crucial for experimental design because they allow researchers to prioritize sgRNAs that maximize both specificity and activity across a variety of genomic settings, as opposed to choosing guides only on the basis of efficiency or PAM availability.

**Figure 7.**
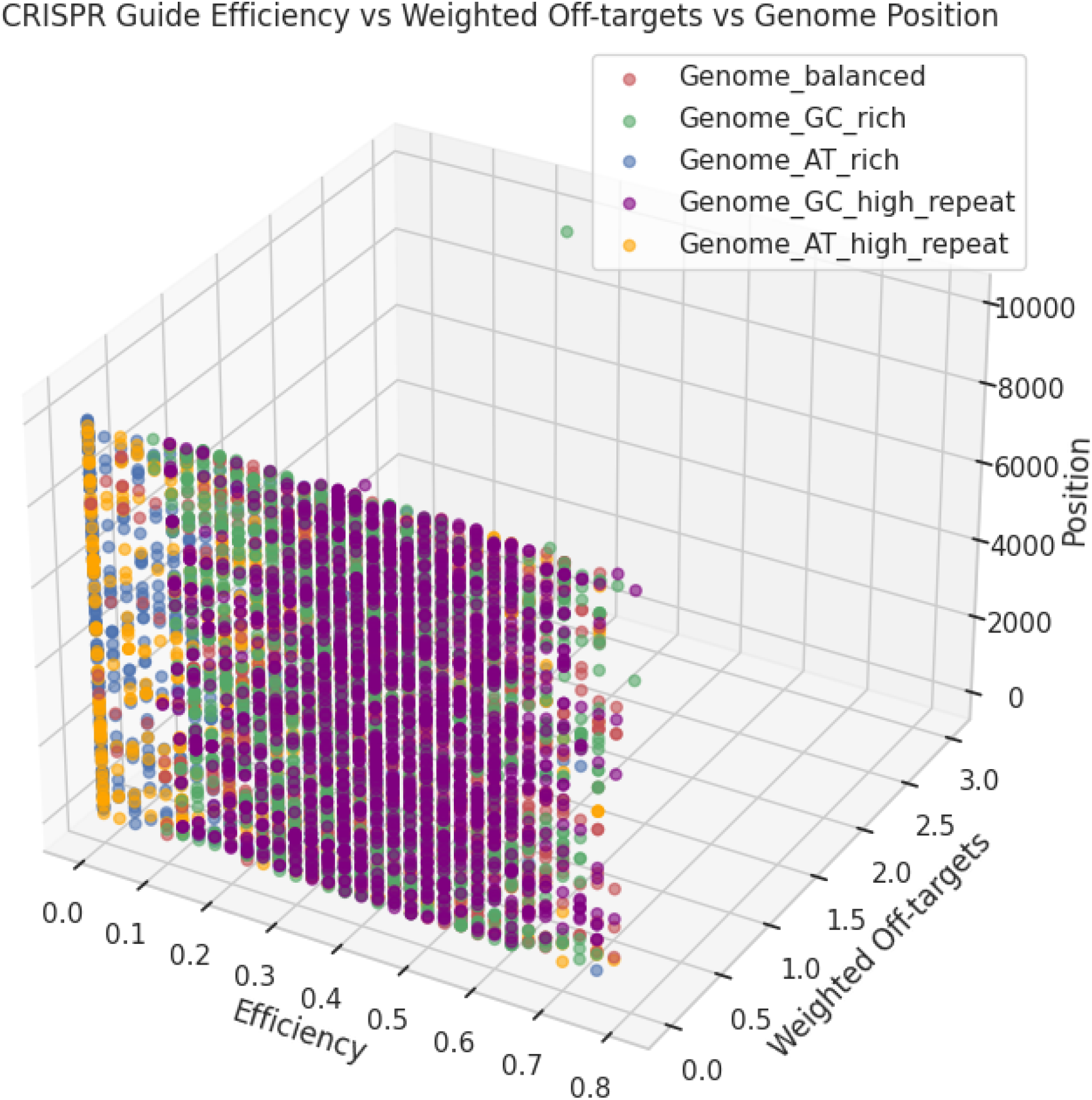

### 3.5 Top-ranked Guides

The top five guidelines for each genome are compiled in Supplementary Table 1. Composite scores above 0.75 and zero off-targets were consistently displayed by high-ranking guides. Multiple guides with maximal composite scores (0.8) were found in the GC-high repeat genome, demonstrating that precision CRISPR targeting is possible even in complicated genomic landscapes. Additionally, the AT-high repeat genome generated highly specific guides, indicating that when employing integrated computational methodologies, optimal guide creation is not impeded by repetitive material. These findings highlight our Python-based pipeline’s translational potential for genome editing research aimed at synthetic constructions or non-model organisms.

### 3.6 Novelty and Biological Relevance

A scalable computational framework for CRISPR guide optimization is established by our analysis, taking into consideration local GC penalties, seed-region mismatch weighting, PAM variability, genome composition, and repeat content. Our pipeline outperforms traditional guide design techniques that frequently overlook genomic variability, repeated regions, or alternative PAMs by incorporating these variables. Additionally, off-target predictions and genome-wide visualizations offer practical insights for experimental design, decreasing trial-and-error and improving the efficacy and safety of genome editing projects. To sum up, this work shows that high-specificity, computationally generated sgRNAs may be consistently found in a variety of genomic settings, offering experimentalists a reliable resource and improving the accuracy of CRISPR-based applications.

## 4. Discussion

The current work integrates weighted off-target scoring, efficiency penalties, and PAM diversity to create a comprehensive, genome composition-aware computational framework for CRISPR/Cas9 guide RNA creation. By specifically taking into account repetitive elements, numerous PAM sequences, and genome-wide sequence composition, our method improves on previous approaches [9,10,18,19] and increases predictive power for both model and non-model organisms.

### Genome Composition Influences Guide Selection

According to our findings, highly repetitive genomes exhibit a minor decrease in maximal composite scores, although GC-rich genomes often favor somewhat better efficiency guides when compared to their AT-rich counterparts. These findings support previous reports that sgRNA activity and specificity are modulated by repeat density, GC content, and local sequence context [31–36,42,43]. We demonstrate that computational guide selection procedures can generalize beyond the human or reference genomes commonly employed in previous studies [37–40] by simulating five synthetic genome types. This offers an essential basis for creating sgRNAs that function well in organisms with unusual genomic structures.

### Weighted Off-target Scoring Improves Predictive Specificity

Our use of seed-region weighted mismatch penalties is consistent with biochemical research showing that cleavage specificity is substantially impacted by mismatches in the first 10 nucleotides next to the PAM [68–70]. The efficiency of integrating positional weighting with genome-wide off-target enumeration was demonstrated by the fact that most top-ranked guides showed zero anticipated off-targets across all genome types. The shortcomings of traditional off-target prediction methods that only use sequence similarity are addressed by our method, which provides a more complex and biologically informed measure of specificity [7,9,14,15,18,19,22].

### Targeting Potential Is Increased by PAM Diversity

Our workflow captures the versatility of many Cas9 orthologs by using alternate PAM sequences (e.g., NGG, NAG) [46–50,73]. Because PAM selection can limit or increase the number of target sites available, this is very important for precision genome editing. Our findings demonstrate that efficiency scores differ depending on the type of PAM, highlighting the need of multi-PAM screening for the best guide selection—a factor that was frequently disregarded in previous computational research [11,12,26].

### Rational Guide Prioritization is Facilitated by Integrative Efficiency-Off-target Landscapes

Clusters of the best-performing guides that strike a balance between specificity and activity are shown by the 3D scatterplots that include efficiency, off-target counts, and genomic location. The conventional trade-off between guide potency and safety, which has hampered experimental CRISPR workflows, is lessened by this dual consideration of efficiency and specificity [4,5,24,25]. Additionally, violin plots and genome-wide heatmaps offer visual aids for locating “safe-harbor” loci, simplifying experimental planning and lowering the need for trial-and-error methods [28,64,65].

Our Python-based workflow guarantees reproducibility, accessibility, and adaptability for a variety of genomic scenarios and is fully compatible with Google Colab [61–65]. The platform may easily support new CRISPR modalities like base editors and Cas12a nucleases by methodically integrating sequence-based scoring with structural, thermodynamic, and positional data [46–50,82–84]. This adaptability bridges a crucial gap between in silico prediction and experimental validation, making our method a useful tool for both editing in poorly described organisms and synthetic genome initiatives. The possibility of precision genome editing across complex genomic landscapes is demonstrated by the discovery of high-specificity guides even in repeat-rich or compositionally extreme areas. By using these insights, researchers can prioritize sgRNAs that minimize the danger of off-target mutagenesis while maximizing both activity and safety [1–6,22,24–27]. Additionally, the pipeline provides mechanistic insights into guide behavior by integrating local GC penalties and repeat-awareness, offering logical criteria for guide selection that go beyond purely empirical techniques [31–36,42–44]. Real genomes include further levels of complexity, such as chromatin accessibility, epigenetic changes, and three-dimensional folding, whereas synthetic genomes offer controlled settings for testing scoring systems [12,16,17,46]. Predictive accuracy will be further improved by future research that takes these characteristics into account and validates them experimentally in a variety of organisms. Furthermore, adding programmable base editors and recently identified Cas variations to the scoring system might improve the tool’s suitability for next-generation genome engineering techniques [83,84]. Lastly, in summary by taking into consideration factors including genome composition, PAM variability, off-target weighing, and efficiency penalties, this work offers a strong, broadly applicable computational paradigm for CRISPR/Cas9 guide RNA design, addressing significant shortcomings of current techniques. Our results give experimentalists practical insights that will speed up the adoption of CRISPR technologies in a wide range of genomic contexts and allow for the logical selection of highly effective, targeted guides.

## Supporting information

Supplementary Codes with Figures

Simulation Top Guides

All Simulation Guides

