## Supplementary Codes with Figures for "Weighted Off-target and Efficiency Scoring Reveal Genome Composition-Dependent Optimal CRISPR/Cas9 Guide Design"

```python
# ===============================
# CRISPR/Cas9 Guide Optimization
# Full upgraded simulation addressing limitations
# Google Colab compatible, Python-only
# ===============================

import numpy as np
import pandas as pd
import matplotlib.pyplot as plt
import seaborn as sns
from mpl_toolkits.mplot3d import Axes3D
import random

sns.set(style="whitegrid")

# ------------------------
# 1️⃣ Function: Simulate genome with adjustable GC content and repeats
# ------------------------
def simulate_genome(length=50000, gc_fraction=0.5, repeat_fraction=0.1, seed=None):
    if seed: np.random.seed(seed)
    # Base composition
    num_gc = int(length * gc_fraction)
    num_at = length - num_gc
    genome_list = ['G','C']*(num_gc//2) + ['A','T']*(num_at//2)
    while len(genome_list) < length:
        genome_list.append(np.random.choice(['A','T','G','C']))
    # Shuffle
    np.random.shuffle(genome_list)
    # Introduce repeats
    num_repeats = int(length * repeat_fraction)
    for _ in range(num_repeats):
        start = np.random.randint(0, length-10)
        repeat_seq = genome_list[start:start+10]
        insert_pos = np.random.randint(0, length-10)
        genome_list[insert_pos:insert_pos+10] = repeat_seq
    return ''.join(genome_list)

# ------------------------
# 2️⃣ Function: Find candidate guides with multiple PAMs
# ------------------------
def find_guides(genome, guide_len=20, pams=['NGG','NAG']):
    guides = []
    for i in range(len(genome)-guide_len-3):
        seq = genome[i:i+guide_len]
        pam_seq = genome[i+guide_len:i+guide_len+3]
        for pam in pams:
            if pam_seq[1:] == pam[1:]:
                guides.append((seq, i, pam_seq))
    return guides

# ------------------------
# 3️⃣ Weighted off-target scoring (seed region more critical)
# ------------------------
def weighted_offtargets(guide, genome, max_mismatches=2, seed_len=10):
    count = 0
    g_len = len(guide)
    for i in range(len(genome)-g_len):
        window = genome[i:i+g_len]
        mismatches = [a!=b for a,b in zip(guide, window)]
        total_mm = sum(mismatches)
        if 0 < total_mm <= max_mismatches:
            # Seed mismatches weighted double
            seed_mm = sum(mismatches[:seed_len])
            count += 1 + seed_mm  # each seed mismatch adds 1 extra penalty
    return count

# ------------------------
# 4️⃣ Efficiency scoring (global + local GC penalties)
# ------------------------
def efficiency_score(guide, window_size=5):
    # Base efficiency
    gc_content = (guide.count('G') + guide.count('C')) / len(guide)
    score = 0.5 + 0.5*gc_content  # 0.5-1
    # Local GC penalty (sliding window)
    penalties = []
    for i in range(len(guide)-window_size+1):
        sub = guide[i:i+window_size]
        sub_gc = (sub.count('G') + sub.count('C'))/window_size
        if sub_gc < 0.3 or sub_gc > 0.7:
            penalties.append(0.05)  # small penalty per window
    score -= sum(penalties)
    [illegible]
```

```
    return max(score,0)  # avoid negative

# ------------------------
# 5️⃣ Analyze genome
# ------------------------
def analyze_genome(genome, genome_id='G1'):
    guides = find_guides(genome)
    results = []
    for g, pos, pam_seq in guides:
        off_targets = weighted_offtargets(g, genome)
        eff = efficiency_score(g)
        comp_score = eff / (1 + off_targets)
        results.append({'genome': genome_id, 'guide': g, 'position': pos,
                        'PAM': pam_seq, 'off_targets': off_targets,
                        'efficiency': eff, 'score': comp_score})
    return pd.DataFrame(results)

# ------------------------
# 6️⃣ Simulate multiple genomes (balanced, GC-rich, AT-rich)
# ------------------------
genomes = {
    'Genome_balanced': simulate_genome(length=10000, gc_fraction=0.5, seed=42),
    'Genome_GC_rich': simulate_genome(length=10000, gc_fraction=0.7, seed=123),
    'Genome_AT_rich': simulate_genome(length=10000, gc_fraction=0.3, seed=999),
    'Genome_GC_high_repeat': simulate_genome(length=10000, gc_fraction=0.65, repeat_fraction=0.2, seed=2025),
    'Genome_AT_high_repeat': simulate_genome(length=10000, gc_fraction=0.35, repeat_fraction=0.2, seed=2026)
}


df_all = pd.concat([analyze_genome(g, gid) for gid, g in genomes.items()], ignore_index=True)

# ------------------------
# 7️⃣ Top guides per genome
# ------------------------
top_guides = df_all.sort_values('score', ascending=False).groupby('genome').head(5)
print("Top 5 guides per genome:")
print(top_guides[['genome','guide','position','PAM','off_targets','efficiency','score']])

# ------------------------
# 8️⃣ Visualizations
# ------------------------

# 8a. Off-target distribution violin plot per genome
plt.figure(figsize=(12,6))
sns.violinplot(x='genome', y='off_targets', data=df_all, inner='quartile', hue='genome', legend=False)
plt.title('Off-target distribution per genome')
plt.ylabel('Weighted Off-targets')
plt.xlabel('Genome')
plt.show()

# 8b. 3D scatter: Efficiency vs Off-target vs Position
fig = plt.figure(figsize=(12,8))
ax = fig.add_subplot(111, projection='3d')
colors = ['r','g','b','purple','orange']
for gid, color in zip(df_all['genome'].unique(), colors):
    df_g = df_all[df_all['genome']==gid]
    ax.scatter(df_g['efficiency'], df_g['off_targets'], df_g['position'], label=gid, alpha=0.6, c=color)
ax.set_xlabel('Efficiency')
ax.set_ylabel('Weighted Off-targets')
ax.set_zlabel('Position')
ax.set_title('CRISPR Guide Efficiency vs Weighted Off-targets vs Genome Position')
ax.legend()
plt.show()

# 8c. Genome-wide heatmap of top guides
for gid in df_all['genome'].unique():
    df_g = df_all[df_all['genome']==gid]
    plt.figure(figsize=(12,2))
    plt.scatter(df_g['position'], [1]*len(df_g), c=df_g['score'], cmap='viridis', s=50)
    plt.colorbar(label='Composite Score')
    plt.title(f'Genome-wide Guide Score Heatmap: {gid}')
    plt.xlabel('Genome Position')
    plt.yticks([])
    plt.show()

# ------------------------
# 9️⃣ Save results
# ------------------------
df_all.to_csv('crispr_full_simulation_all_guides.csv', index=False)
top_guides.to_csv('crispr_full_simulation_top_guides.csv', index=False)
print("All results saved to 'crispr_full_simulation_all_guides.csv' and 'crispr_full_simulation_top_guides.csv'")
```

```
Top 5 guides per genome:
                   genome                 guide  position  PAM  off_targets  \
2673         Genome_GC_rich  CCCATCCTGACCATCACCAG      7478  AGG            0
5592  Genome_GC_high_repeat  GGTGTCCTGTCCTCTCAGTG      9613  CGG            0
153         Genome_balanced  GCATCCTTCTCGTCTGGATG      1309  TAG            0
3137         Genome_GC_rich  GTCAGTCCTCTCCAACTGAC      9602  AGG            0
756         Genome_balanced  CCTCACGTAGACTTGCCAAG      6076  TGG            0
755         Genome_balanced  GTACCTCACGTAGACTTGCC      6073  AAG            0
61          Genome_balanced  GCCTAACGAGTGCTTCGAGA       551  TGG            0
5706  Genome_AT_high_repeat  AGGTGAGAGTCTGCAGTAGC       480  TGG            0
280         Genome_balanced  CCAGTGGAAGACGTATGCCA      2263  TGG            0
4791  Genome_GC_high_repeat  CGTCATGACGAGTCAGTCGT      5090  CGG            0
4170  Genome_GC_high_repeat  GCTACACCTTGTCCACTTGG      1512  GAG            0
5027  Genome_GC_high_repeat  CGCTACACTTGGGTTCACAG      6415  TAG            0
5640  Genome_GC_high_repeat  CGGTTCCACGAGTGGATCCC      9881  GAG            0
2166         Genome_GC_rich  CGCTTCGCTACCTCCAGTCT      4659  CGG            0
2164         Genome_GC_rich  GGCACTCTGAGAGTCAGAGC      4626  GGG            0
2222         Genome_GC_rich  CGGATGCGTACATCGCATCG      5061  CGG            0
3218         Genome_AT_rich  CAACGATAGCAGACAGTTCG        75  TAG            0
5984  Genome_AT_high_repeat  TGGTACGCAACCTTGTCGTA      4057  AGG            0
5985  Genome_AT_high_repeat  GGTACGCAACCTTGTCGTAA      4058  GGG            0
5983  Genome_AT_high_repeat  TTGGTACGCAACCTTGTCGT      4056  AAG            0
6154  Genome_AT_high_repeat  TGACTACTGACACCTTAGGG      6220  CGG            0
3588         Genome_AT_rich  ATTGCAGAGAACGTCTGACA      5031  TAG            0
3589         Genome_AT_rich  GCAGAGAACGTCTGACATAG      5034  TAG            0
3632         Genome_AT_rich  ACTACCCAACACAAAGGGAT      5579  AAG            0
3693         Genome_AT_rich  TTTGCAGAAGCCTTTCTCGT      6349  AAG            0

      efficiency  score
2673       0.800  0.800
5592       0.800  0.800
153        0.775  0.775
3137       0.775  0.775
756        0.775  0.775
755        0.775  0.775
61         0.775  0.775
5706       0.775  0.775
280        0.775  0.775
4791       0.775  0.775
4170       0.775  0.775
5027       0.775  0.775
5640       0.775  0.775
2166       0.750  0.750
2164       0.750  0.750
2222       0.750  0.750
3218       0.750  0.750
5984       0.750  0.750
5985       0.750  0.750
5983       0.750  0.750
6154       0.750  0.750
3588       0.725  0.725
3589       0.700  0.700
3632       0.675  0.675
3693       0.675  0.675
```

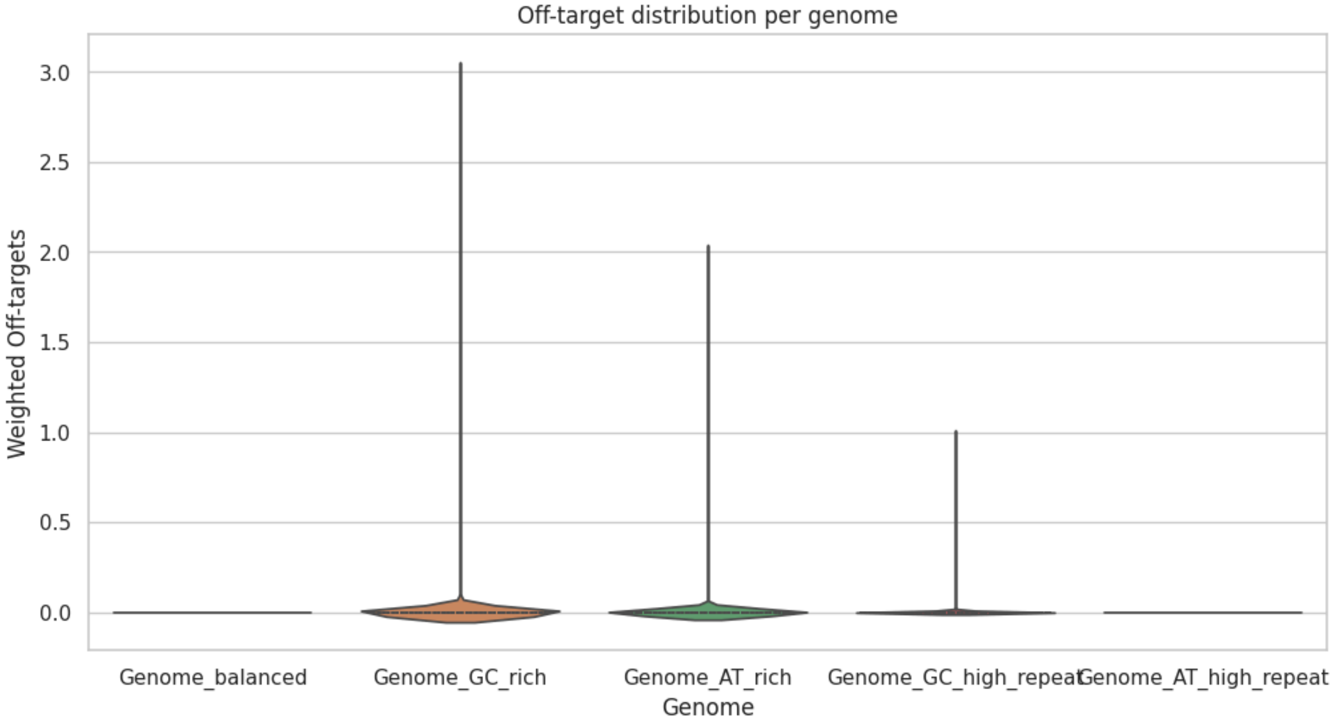


CRISPR Guide Efficiency vs Weighted Off-targets vs Genome Position

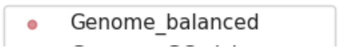

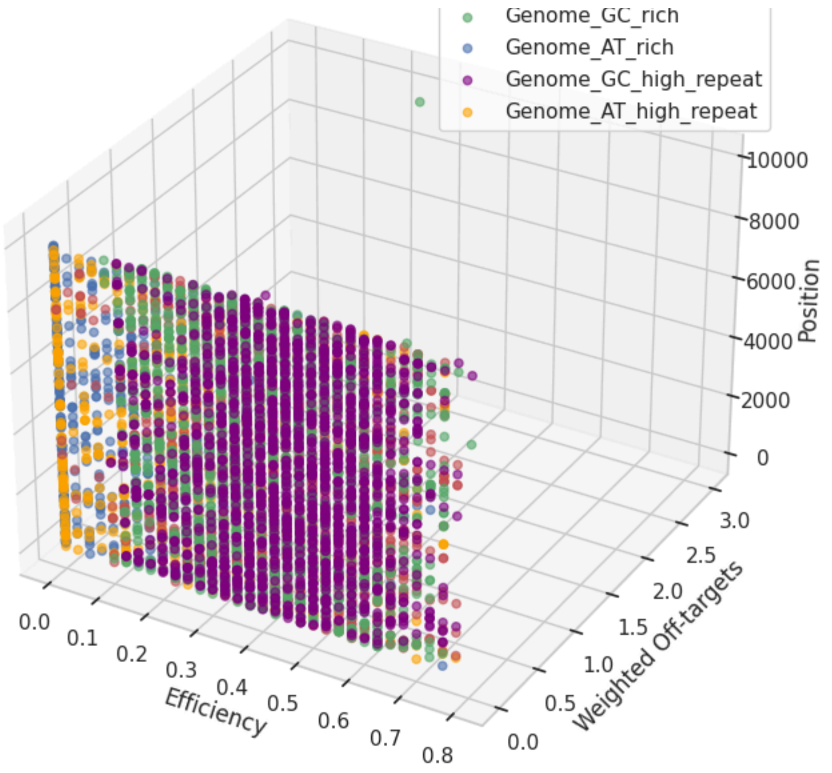
Genome_GC_rich
Genome_AT_rich
Genome_GC_high_repeat
Genome_AT_high_repeat
Efficiency
Weighted Off-targets
Position

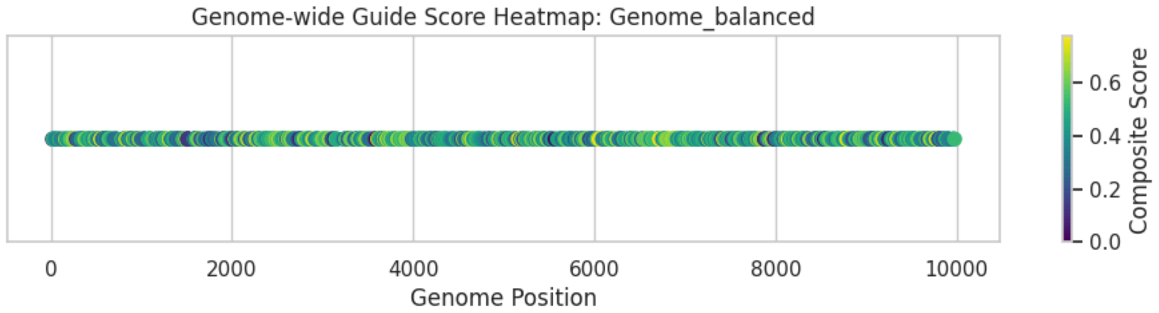
Genome-wide Guide Score Heatmap: Genome_balanced
Genome Position
Composite Score

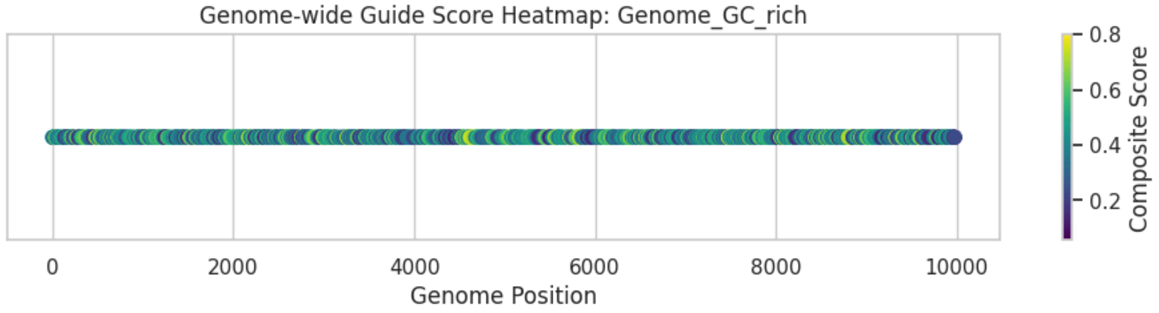
Genome-wide Guide Score Heatmap: Genome_GC_rich
Genome Position
Composite Score

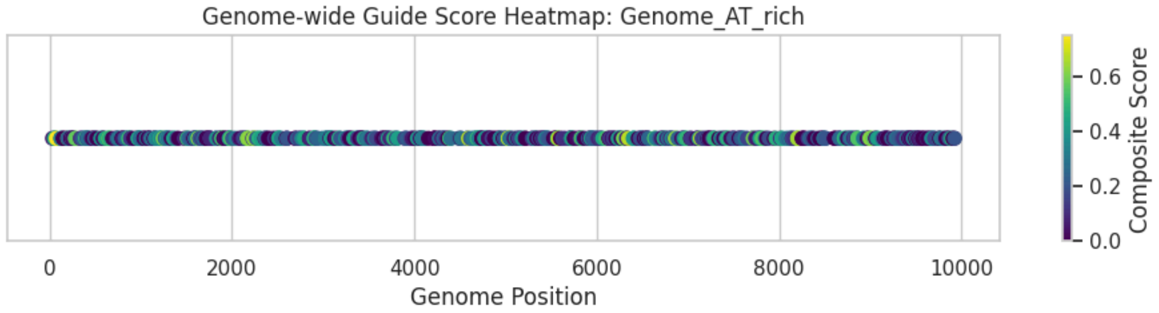
Genome-wide Guide Score Heatmap: Genome_AT_rich
Genome Position
Composite Score

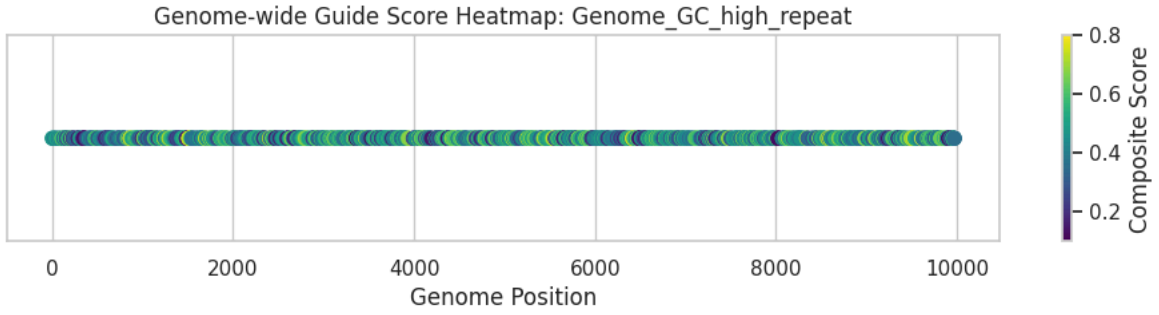
Genome-wide Guide Score Heatmap: Genome_GC_high_repeat
Genome Position
Composite Score
